# Microarray Analysis of Potential Biomarkers for brachial plexus avulsion caused neuropathic pain

**DOI:** 10.1101/2021.08.05.455327

**Authors:** Le Wang, Jie Lao

## Abstract

Nerve injury-induced neuropathic pain remains a challenging clinical problem due to a lack of satisfactory treatment. Pain after BPA (Brachial Plexus Avulsion) is resistant to most traditional pain relief treatments due to the lack of understanding of the cellular or molecular mechanism of pain development. The present study aimed to investigate the expression of mRNA in the brachial plexus avulsion neuropathic pain model and analyze biological functions. Sprague-Dawley rats were treated with complete brachial plexus avulsion. An animal behavior test was carried out to distinguish the pain group from the control group. In this study, a microarray mRNA assay and reverse transcriptase quantitative polymerase chain reaction (RT-PCR) was conducted. The whole blood was collected from two groups for Microarray mRNA analysis. The predicted mRNA targets were studied by gene ontology analysis and pathway analysis. The PIK3CB, HRAS, and JUN genes were verified by RT-PCR. In total, differentially expressed genes(DEGs) were identified between individuals with or without neuropathic pain (case and control), and A biological processes were enriched. We identified 3 targeted mRNAs, including PIK3CB, HRAS, and JUN, which may be potential biomarkers for BPA-caused NP. The results showed that PIK3CB, HRAS, and JUN gene expression was increased in the control group but decreased in the neuropathic pain group. The PIK3CB gene was part of the Neurotrophin signaling pathway. The function of the HRAS gene was synergetic in the aspect of axon guidance and the Neurotrophin signaling pathway. The JUN gene participates in axon regeneration. These results suggest that PIK3CB, HRAS, and JUN genes might become potential biomarkers for the prediction of and new targets for the prevention and treatment of neuropathic pain after BPA. These findings indicate that mRNA expression changes in the blood may play an important role in the development of NP after BPA, which is of theoretical and clinical importance for future research and clinical-treatment strategies.

## 1. Introduction

Nerve injury-induced neuropathic pain remains an intractable disease due to a lack of satisfactory treatment [9]. In 2017, Palma Ciaramitaro et al. [1] investigate the prevalence of neuropathic pain after traumatic brachial plexus injury. Of the 107 patients enrolled, 69% had neuropathic pain. Neuropathic pain can significantly impair function, appetite, sleep, mood, and quality of life. Brachial plexus avulsion (BPA) induces a characteristic of pain are allodynia, hyperalgesia, and persistent pain, which is often difficult to cure [10]. The pain may be manifested as burning or pressure. Pain after BPA is resistant to most pain relief treatments the exact molecular mechanisms responsible for this pathology remain unknown[3]. Previous studies demonstrated that the mRNA plays a key role in the development and maintenance of neuropathic pain [11–13].

Some authors proved that c-Jun plays a vital role in the survival of ventral horn motoneurons in adult mice[4]. At present, there is no literature to prove that the HRAS gene is related to neuropathic pain caused by brachial plexus injury. Previous studies have shown that PIK3CB influences the early development of neuropathy in sensory neurons[2]. The spinal cord plays an important role in the process of central sensitization [14]. Furthermore, we aimed to investigating the mRNA changes of neuropathic pain caused by the brachial plexus avulsion model, thereby providing a novel insight into the mechanism of neuropathic pain.

## 2. Material and methods

### 2.1 Animals

This study was carried out in strict accordance with the recommendations in the Guide for the Care and Use of Laboratory Animals of the National Institutes of Health. The protocol was approved by the Committee on the Ethics of Animal Experiments of the University of Fudan (GB/T 35892-2018). All surgery was performed under sodium pentobarbital anesthesia, and all efforts were made to minimize suffering. The experiments were conducted in male Sprague- Dawley rats (n=20, age, eight weeks; weight, 200- 250g; supplied by the Department of Laboratory Animal Science, Fudan University, Shanghai, China).

### 2.2 Surgery procedure

All surgical procedures were performed after anesthesia induced by a 1% sodium pentobarbital solution (40 mg/kg body weight). Place the rat prone on a sterilized pad with the head oriented away from the surgeon and the right forepaw abducted and extended. Use the fingertips to locate the clavicle. With a scalpel, make a 1.5 cm horizontal incision in the skin under the clavicle 2 mm. Use micro-dissecting scissors to separate the skin from the superficial fascia, exposing the pectoralis major muscle. The pectoralis major muscle was cut paralleled with the muscle fibers to expose the brachial plexus, leaving the cephalic vein intact. The subclavian vessels were located and the upper, middle and lower trunks were dissected. In the complete brachial plexus avulsion (BPA) group (n = 20), the upper, middle, and lower trunk were grasped with forceps and haul out from the spinal cord. The tissue layers were then brought together, and the skin was closed with 4–0 silk sutures (Ethicon), as described previously[7].

### 2.3 Animal pain tests

#### Mechanical allodynia

Mechanical allodynia was assessed by using the von Frey filaments (Stoelting, USA; bending force: 2.0, 4.0, 6.0, 8.0, 10.0, 15.0, and 26.0 g). The filaments were applied to the left forepaw. The threshold was the lowest force that evoked a withdrawal response. Each filament was applied five times. When rats showed at least two withdrawal responses to a filament, the bending force of the filament was defined as the withdrawal threshold.

#### Cold allodynia

Cold allodynia was assessed by an acetone spray test as described by Choi et al. [8]. 250 μl acetone was squirted onto the surface of the paw. Neuropathy rats frequently responded with a withdrawal that was clearly exaggerated in amplitude and duration. The withdrawal responses were assessed on a scale of 3-0 points: 3 points, a vigorous response in which the rat licked the paw; 2 points, a response in which the paw has elevated the paw; 1 point, a response in which the paw had little or no weight born on it and 0 points, the paw was not moved[5].

### 2.4 mRNA microarray

The whole blood was collected from the rats. Total RNA was extracted from whole blood using a QIAamp RNA blood mini kit (Qiagen) per the manufacture’s instruction. Purified total RNA for each strain used in Affymetrix GeneChip assays (Affymetrix GeneChip Rat Gene 1.0). Microarrays were processed using an Agilent GeneArray Scanner with Affymetrix Microarray Suite version 5.0.0.032 software.

### 2.5 Reverse transcription-quantitative polymerase chain reaction (RT-PCR) assay

RNA was reverse transcribed into cDNA using Takara PrimeScript RT master mix (RR036A; Takara Biotechnology Co., Ltd., Dalian China). RT-qPCR was performed using an ABI StepOne Plus Real-Time PCR system (Thermo Fisher Scientific, Inc.) and SYBR Premix Ex Taq II master mix (Takara Biotechnology Co., Ltd.) according to the manufacturer’s protocol. The reaction system (10 μl) consisted of cDNA (1 μl), forward primers (10 μM; 0.2 μl), reverse primers (10 μM; 0.2 μl), ROX reference dye (0.2 μl), RNase-free water (3.4 μl), and SYBR-Green mixture (5 μl). The thermocycling conditions were as follows: Initial denaturation, 95°C for 30 sec, followed by 40 cycles of 95°C for 5 sec and 60°C for 30 sec. Rat actin was used as a housekeeping gene. The relative expression of genes was calculated using the 2^−ΔΔCt^ method.

### 2.6 Bioinformatic evaluation

GO analysis was applied to analyze the function of the expression genes according to the Gene Ontology, which is the crucial function of NCBI that can organize genes into hierarchical classification and uncover the gene network on the basis of biological process and molecular function.

Pathway analysis was applied to find out the significant pathway of the differential genes according to KEGG, Biocarta, and Reactome. Still, we turn to Fisher’s exact test and *χ*^2^ test to select the most significant pathway, and the threshold of significance was decided by *P*-value and FDR. The enrichment Re was calculated like the equation above.

### 2.7 Statistical analysis

The random variance model t‑test was adopted to filter the differinitially expressed mRNAs between the control and pain groups using GraphPad 5.0. Following the significance analysis and false discovery rate analysis, differentially expressed genes were selected according to their P-values. P<0.05 was considered to indicate a statistically significant difference.

## 3. Results

### mRNA microarray

Animals exhibiting significant decreases in the pain threshold(mechanical threshold decreases from 15 g pre-surgery to 8 g post-surgery and allodynia score increases from 0 pre-surgery to 2-3 post-surgery) were placed in the NP(Neuropathic Pain) group. There were 10 rats do the BPA surgery and 6 rats had neuropathic pain. There were 6 rats in the NP group. The sham-operated animals whose brachial plexus was just dissected but not used were assigned to the control group. There were 10 rats in the control in the control group.

To functionally investigate a possible link between mRNA expression and the brachial plexus injury neuropathic pain, the differential expression of mRNA in the neuropathic pain and control group was analyzed. The whole blood was harvest from the rat after 2 weeks. The expression of 2717 mRNAs was detected between the pain and control group according to the changes: down and up. By contrast to the control group and the pain group, 1154 mRNAs exhibited decreased expression, and 1563 mRNAs exhibited increased expression. The most significant top 10 upregulated or downregulated genes are shown in Table 1.

**Table1.** The most significant Upregulated Genes or Downregulated Genes in neuropathic pain group

| Probe set ID | Gene symbol | Gene description | P-value | Trend |
| --- | --- | --- | --- | --- |
| 10937619 | LOC685774 | Hypothetical protein | 0.0253862 | down |
| 10937311 | Mir448 | microRNA | 0.0380793 | down |
| 10936853 | Midlip1 | MID1 protein | 0.0390031 | down |
| 10934445 | Ogt | GlcNAc transferase | 0.0395949 | down |
| 10929600 | Pde6d | Phosphodiesterase 6D | 0.0389264 | down |
| 10929445 | Tm4sf20 | transmembrane | 0.0374896 | down |
| 10925373 | Ube2f | Ubiquitin-conjugating enzyme E2F | 0.0308392 | down |
| 10911048 | Dapk2 | Death-associated kinase 2 | 0.0257355 | down |
| 10908788 | Zbtb44 | Zinc finger and BTB domain containing 44 | 0.0406164 | down |
| 10905558 | Rpl26 | Ribosomal protein L26 | 0.0278983 | down |
| 10714907 | Ifit1 | Interferon-induced | <1e-07 | up |
| 10886573 | Ifi27 | Interferon, alpha-inducible protein 27 | <1e-07 | up |
| 10811177 | Ctrb1 | Chymotrysinogen B1 | <1e-07 | up |
| 10882317 | Isg15 | ISG 15 Ubiquitin-like modifier | <1e-07 | up |
| 10827820 | RT1-T24-4 | RT1 class I, locus T24, gene 4 | <1e-07 | up |
| 10737262 | Supt4h1 | Suppressor of Ty4 homolog 1 | <1e-07 | up |
| 10732592 | Nprls | Nitrogen permease regulator-like 3 | <1e-07 | up |
| 10785144 | Xpo07 | Exportin7 | <1e-07 | up |
| 10749495 | Lgals3bp | Lectin, galactoside-binding, soluble, 3 binding protein | <1e-07 | up |
| 10704505 | S1c1a5 | Solute carrier family 1(neutral amino acid transporter), member 5 | <1e-07 | up |

We have found 621 GO terms with the P-value <0.05. The top 20 GO terms, ranked by P-value, were shown in table 2. Most of the enriched terms were about inflammatory processes involved in protein modification and regulation of biological processes. The result was similar to the GO analysis.

**Table 2.** The Top 20 most significant GO terms in neuropathic pain group

| GO Terms | GO name | Path-count | Enrichment | Trend |
| --- | --- | --- | --- | --- |
| 0009615 | Response to virus | 84 | 12.30162448 | up |
| 0045087 | Innate immune response | 93 | 10.69962082 | up |
| 0008150 | Biological process | 1408 | 2.582253877 | up |
| 0043066 | Negative regulation of apoptotic | 478 | 4.003318056 | up |
| 0042493 | Response to drug | 462 | 3.893443439 | up |
| 0051607 | Defense response to virus | 96 | 8.371938884 | up |
| 0014070 | Response to an organic cyclic compound | 230 | 5.158362344 | up |
| 0008285 | Negative regulation of cell proliferation | 308 | 4.349059161 | up |
| 0032355 | Response to estradiol stimulus | 143 | 6.155591427 | up |
| 0006954 | Inflammatory response | 189 | 5.264892783 | up |
| 0008150 | Biological process | 1408 | 2.44882276 | down |
| 0006355 | Regulation of transcription DNA-dependent | 681 | 2.8479700082 | down |
| 0006351 | Transcription, DNA-dependent | 640 | 2.862061601 | down |
| 0006886 | Intracellular protein transport | 167 | 4.838991083 | down |
| 0015031 | Protein transport | 271 | 3.777150973 | down |
| 0006412 | Translation | 384 | 3.226834158 | down |
| 0045944 | Positive regulation of transcription from RNA polymerase II promoter | 715 | 2.411148564 | down |
| 0000122 | Negative regulation of transcription from RNA polymerase II promoter | 499 | 2.699103242 | down |
| 0015986 | ATP synthesis coupled proton transport | 19 | 14.17739493 | down |
| 006302 | Double-strand break repair | 48 | 7.856639688 | down |

GO: Gene Ontology; Count: enriched gene numbers in each term.

To further investigate the functions of DEGs, we did a KEGG pathway analysis. The top 20 pathways were shown in table 3. The most significantly enriched pathways were Metabolic pathways, antigen processing and presentation, and Herpes simplex infection.

**Table 3.** The Top 20 most significant enriched KEGG pathways

| KEGG Term | Pathname | Path<br>gene<br>count | Enrichment | Trend |
| --- | --- | --- | --- | --- |
| 04145 | Phagosome | 196 | 7.810555227 | up |
| 05168 | Herpes simplex infection | 218 | 6.320100652 | up |
| 01100 | Metabolic pathways | 1272 | 2.768080422 | up |
| 04612 | Antigen processing and presentation | 98 | 9.372666273 | up |
| 05203 | Viral carcinogenesis | 239 | 5.284379834 | up |
| 05169 | Epstein-Barr virus infection | 232 | 5.278858016 | up |
| 04062 | Chemokine signaling pathway | 180 | 5.953378762 | up |
| 04670 | Leukocyte transendothelial migration | 119 | 7.075444147 | up |
| 04516 | Viral myocarditis | 110 | 7.30641939 | up |
| 04144 | Endocytosis | 236 | 4.702880923 | up |
| 00190 | Oxidative phosphorylation | 162 | 7.981348255 | down |
| 05010 | Alzheimer's disease | 214 | 6.545451489 | down |
| 05012 | Parkinson's disease | 164 | 6.898512897 | down |
| 01100 | Metabolic pathways | 1272 | 2.625938872 | down |
| 05016 | Huntingtin's disease | 219 | 4.920009198 | down |
| 04120 | Ubiquitin mediated<br>proteolysis | 136 | 5.149730216 | down |
| 04141 | Protein processing in the<br>endoplasmic reticulum | 165 | 4.571135819 | down |
| 04110 | Cell cycle | 126 | 5.130866735 | down |
| 05168 | Herpes simplex infection | 218 | 3.706933536 | down |
| 05164 | Influenza A | 177 | 3.956854855 | down |

### PCR verification

We aimed to explore the mechanism of neuropathic pain after brachial plexus avulsion and find nerve-related mRNAs. Therefore, we mainly focused on the differentially expressed genes(DEGs) dysregulated only in the neuropathic pain groups. The three nerve-related mRNAs (Pik3cb, Hras, and Jun genes) exhibited decreased expression in the neuropathic pain group.

To validate the microarray results, RT-qPCR was performed for Pik3cb, Hras, and Jun genes. It was found that the relative expression of 3 mRNA among them was significantly altered, which coincided with the results of the microarray (Figure 1).

**Figure 1:**
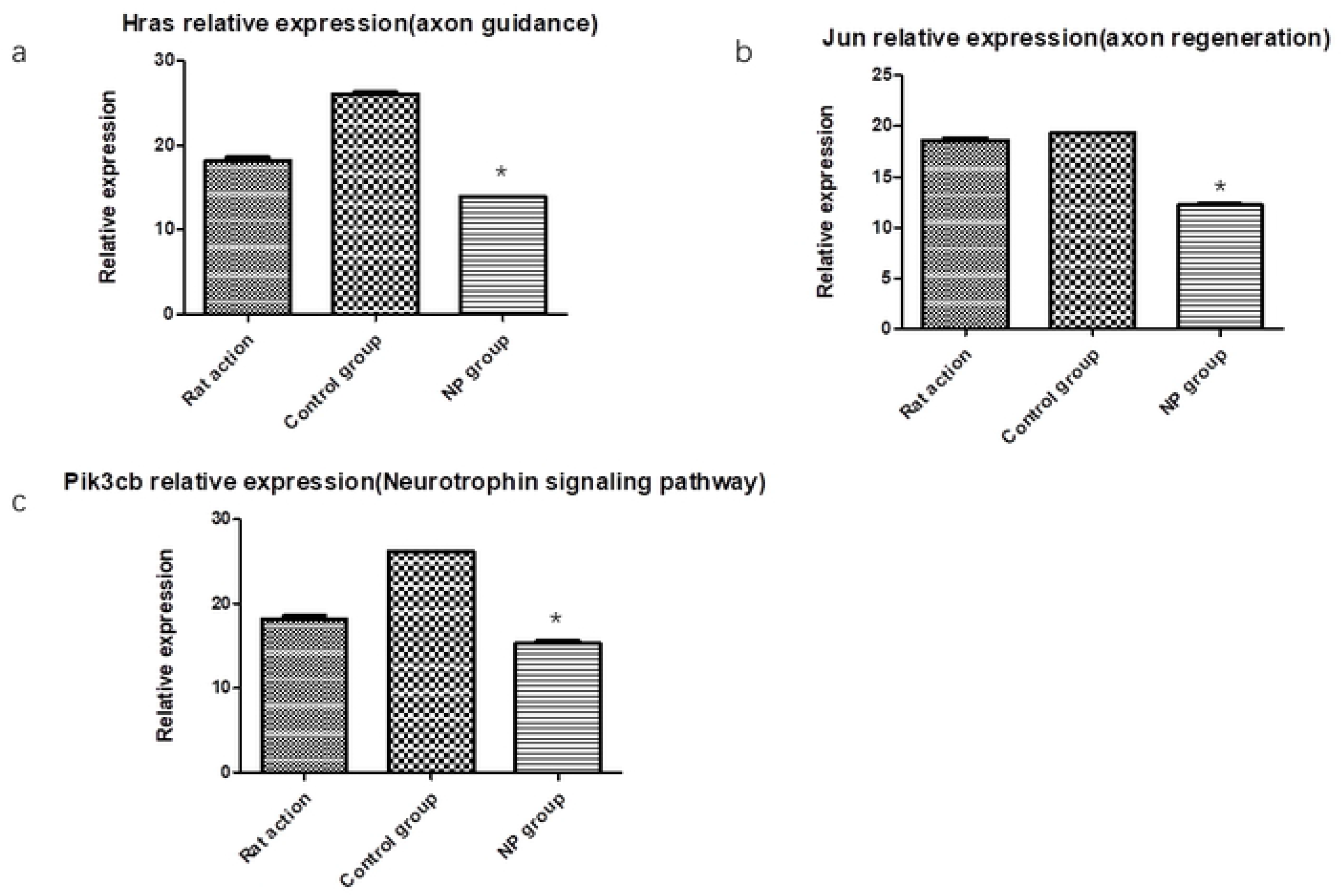
Relative expression of differentially expressed mRNA in rat whole blood in the microarray. (a) Hras were significantly down-regulated in neuropathic pain group versus the control group after 2 weeks. (b) Jun was significantly down-regulated in neuropathic pain group versus the control group after 2 weeks. (c) Pik3cb were significantly down-regulated in neuropathic pain group versus the control group after 2 weeks. Data are presented as mean±SE, * p<0.05. NP group: Neuropathic Pain group

### Bioinformatics analysis of the diff-reg mRNA

The Pik3cb, Hras, and Jun genes were intersection genes, which were involved in neuropathic pain according to GO and pathway analyses. The results showed that Pik3cb, Hras, and Jun gene expression was high in the control group but was low in the neuropathic pain group. The function of the Hras gene was synergetic in the aspect of axon guidance and the Neurotrophin signaling pathway. The Jun gene function was axon regeneration. The low expression of two genes in the neuropathic pain group was revealed that neuropathic pain is unfavorable for nerve regeneration.

## 4. Discussion

Brachial Plexus Avulsion(BPA) has been demonstrated to be a polygenic disease and its pathogenic mechanism is associated with changes in many genes. In this study, we have used microarray to identify differentially expressed genes(DEGs) and activated signaling pathways in association with BPA-induced neuropathic pain(NP) in a rat BPA model. We showed that Jun, HRAS, and PIK3B were the nerve-related downregulated DEG.

Our results are consistent with that of previous studies[7]. Although the precise roles of the three marker genes in BPA-induced NP are not completely understood, our data highlighted the diagnostic and treatment potential of this disease.

It will be very interesting to further this study into BPA patients. Microarray technology can be reliable and useful for identifying novel targets for clinical diagnostic and therapeutic approaches. This technology can be used in pancreatic cancer and renal clear cell carcinoma for diagnosis and effective therapy[15,17].

We found that three genes expressed decreased and were related to nerve regeneration. Some authors proved that the downregulation of c-Jun gene expression is not conducive to the survival of motoneurons. HRAS might serve specific roles in the development and maintenance of nervous tissues[6]. In our study, the Metabolic signaling pathway and Phagosome signaling pathway are involved in BPA, which play a very important role in BPA-induced NP. In the peripheral nervous system, recent studies suggested that the nerve-related gene plays an important role in neuropathic pain after spinal cord injury[16]. Some authors suggested that there is a high possibility of neuropathic pain caused by nerve damage[18,19]. The transcriptome changed play an important role in neuropathic pain[13]. So our research is meaningful and feasible. Ji-An Yang et al[20] proved that Jun is a potential indicator for neuropathic pain. Despite increasing knowledge and ongoing study, the precise molecular mechanisms of neuropathic pain caused by brachial plexus injury remain largely unknown. Numerous studies show a significant modification of gene expression as a consequence of nerve injury. A study by Timo et al reported that miRNAs-494, −720,−690, and −668 showed the highest signal intensities in the rat spinal cord[21]. The exosomes with Ccl3 can be efficiently detected in peripheral blood. Guan Zhang et al proved that Ccl3 can be used as a potential prognostic target for the diagnosis and treatment of spinal cord injury-induced chronic neuropathic pain in clinical applications. The microarray analysis of DEGs and pathway indifferent section by GO and KEGG suggests another method and strategy research the target gene and pathway of nerve-related disease[22].

In summary, our studies indicated that Jun, HRAS, and PIK3B might serve a significant role in neuropathic pain and nerve regeneration. we demonstrated that microarray. The three nerve-related genes were downregulated in the spinal cord in NP rats after brachial plexus avulsion. Furthermore, KEGG analysis found that Metabolic pathways with significance were identified. These results strongly suggest that neuropathic pain may attenuate nerve regeneration via inhibition of neurotrophin signaling pathway and axon guidance pathway, which is of theoretical and clinical importance for future research and clinical-treatment strategies. Several limitations should be acknowledged in our study. First, the sample size was relatively small. Besides, the results were all base on a rat model. In the future, we will perform some more in-depth studies around nerverelated genes.

## Supporting information

**Table1. The most significant Upregulated Genes or Downregulated Genes in neuropathic pain group**

(DOCX)

**Table 2. The Top 20 most significant GO terms in neuropathic pain group**

(DOCX)

**Table 3. The Top 20 most significant enriched KEGG pathways**

(DOCX)

## Acknowledgments

This work was supported by the Ministry of Science and The technology of China (973 Program Grant 2014CB542204).

## Author Contributions

**Conceptualization:** Jie Lao, Le wang.

**Data curation:** Le wang

**Formal analysis:** Jie Lao, Le wang.

**Funding acquisition:** Jie Lao

**Investigation:** Jie Lao, Le wang.

**Methodology:** Jie Lao, Le wang.

**Project administration:** Jie Lao, Le wang.

**Resources:** Jie Lao, Le wang.

**Supervision:** Jie Lao

**Validation:** Le wang.

**Writing – original draft:** Le wang.

**Writing – review & editing:** Jie Lao.

